# Identification of conserved skeletal enhancers associated with craniosynostosis risk genes

**DOI:** 10.1101/2022.09.01.506150

**Authors:** Xuan Anita He, Anna Berenson, Michelle Bernard, Chris Weber, Juan I. Fuxman Bass, Shannon Fisher

## Abstract

Craniosynostosis (CS) is a common congenital defect affecting more than 1/2000 infants. Infants with CS have a premature fusion of one or multiple cranial sutures resulting in restricted brain expansion. Single gene mutations account for 15-20% of cases, largely as part of a syndrome, but the majority are nonsyndromic with complex underlying genetics. Two noncoding genomic regions contributing to CS risk were previously identified by GWAS, one near *BMP2* and one within *BBS9*. We hypothesized that the region within *BBS9* contains distal regulatory elements controlling the neighboring gene encoding BMPER, a secreted modulator of BMP signaling. To identify regulatory sequences that might underlie disease risk, we surveyed conserved noncoding sequences from both risk loci identified from the GWAS for enhancer activity in transgenic *Danio rerio*. We identified enhancers from both regions that direct expression to skeletal tissues, consistent with the endogenous gene expression. Importantly, for each locus, we found a skeletal enhancer that also contains a sequence variant associated with CS risk. We examined the activity of each enhancer during craniofacial development and found that the *BMPER*-associated enhancer is active in the restricted region of cartilage closely associated with frontal bone initiation. We used an enhanced yeast one-hybrid assay to identify transcription factor interactions with several identified enhancers, implicating multiple signaling pathways in their regulation. In a targeted screen focused on risk-associated SNPs, we further identified differential binding to alternate and reference alleles. Additionally, we found that the risk allele of the *BMPER* enhancer directs significantly broader expression than the reference allele in transgenic zebrafish. Our findings support a specific genetic mechanism to explain the contribution of two risk loci to CS. More broadly, our combined *in vivo* approach is applicable to many complex genetic diseases to build a link between association studies and specific genetic mechanisms.

**Author Summary:** Genome–wide association studies (GWASs) provide a wealth of information implicating regions of the genome in disease risk. The great challenge is linking those regions to specific genetic mechanisms. We used complementary approaches in zebrafish and yeast to evaluate the genetic risk of craniosynostosis (CS), a craniofacial birth defect affecting 1/2000 infants where two or more skull bones are fused prematurely. Using transgenic zebrafish, we identified sequences regulating expression of two genes in the BMP signaling pathway that had been previously implicated by GWAS. These included one from each region containing a sequence variant linked to disease risk. We used an assay in cultured yeast to detect proteins binding to identified DNA sequences that could alter expression of the target genes, including changes in protein binding caused by the sequence variants. Finally, we found that transgenic fish carrying one of the variant sequences showed broader and more sustained activity throughout the skeleton. Taken together, our results support a model where variant sequences lead to increased gene expression and BMP pathway activity, contributing to aberrant skull growth in CS. Importantly, our paradigm is broadly applicable to other complex genetic diseases, potentially illuminating many connections between genome variation and disease risk.

## Introduction

The bone morphogenetic protein (BMP) pathway is subject to complex regulation through a network of extracellular factors (1). Many of these are required in the development of multiple organ systems and are implicated in human disease, including several skeletal conditions and some cancers (2). BMP signaling plays a particular role in the formation of the skeletal system, and recent findings have implicated dysregulation of the pathway in the craniofacial defect craniosynostosis (CS), in which one or more cranial sutures fuse prematurely. CS is one of the most common structural birth defects, affecting ~1/2000 infants. Mutations in single genes account for only 15-20% of CS cases, which are part of syndromes of coincident defects and mostly affect the coronal sutures. The genetic causes of the remaining cases, which largely affect the midline sagittal and metopic sutures, are more complex (3). An association study focused on nonsyndromic CS (NCS) identified two risk regions, one downstream of *BMP2* (hg38, chr20: 7112785-7245836, and the other in an intron of *BBS9* (hg38, chr7: 33179156-33384149) (4). There were no coding sequence variants associated with either risk allele, so the causal mutations are presumed to affect noncoding sequences. However, in the absence of functional data, it is difficult to predict which of the several linked single nucleotide polymorphisms (SNPs) in each risk locus might contribute to disease risk or to explore the mechanism further.

*BBS9* is one of 14 genes associated with Bardet Biedel syndrome (BBS) and encoding components of functional cilia (5). While BMP2 is one of the most osteogenic BMP ligands(6), BBS9 has no known function in osteogenesis, and CS is not a consistent feature of BBS. However, the adjacent gene encodes Bone Morphogenetic Protein Binding Endothelial Regulator (BMPER), an extracellular regulator of BMP signaling. Homologous to the fly *crossveinless*, the vertebrate gene was originally described as a negative regulator of BMP signaling during early endothelial cell differentiation (7). Later studies supported both pro-and anti-BMP activities (8) (9), suggesting the role of BMPER is context-dependent.

Several lines of evidence suggest that BMPER activity is largely pro-BMP in osteogenesis. Upregulation of *BMPER* promotes BMP2-induced osteogenic differentiation in human bone mesenchymal stem cells (10). Homozygous null mutations of *BMPER* cause diaphanospondylodysostosis (DSD), a lethal perinatal skeletal condition characterized by reduced skeletal structures (11) (12). *BMPER* mutations that result in a truncated protein cause a less severe form of DSD (13). *Bmper* null mutant mice display similar skeletal defects (14). Although the expression of the mammalian gene has not been described during skeletal development, zebrafish *bmper* (previously named *crossveinless 2)* is prominently expressed in cranial bones and cartilages in larvae (8) (15) and adult fish (16). Taken together, the mammalian mutant phenotypes and the expression of the zebrafish gene in developing bone support a conserved role of BMPER in skeletal development across vertebrates. In addition, it is plausible that the two genes comprise a conserved regulatory domain, given 1) the syntenic relationship of *BBS9* and *BMPER* is conserved from mammals to zebrafish (17); 2) the two genes are in the same topologically associated domains (TADs) (18).

We hypothesized that both regions implicated in CS risk harbored noncoding sequences important in regulating genes in the BMP signaling pathway, *BMPER*, and *BMP2*. Given that *BBS9* and *BMPER* are in the same TAD, we also predicted the existence of a broader regulatory domain in the intergenic region. We therefore aimed to identify enhancers regulating expression of both *BMP2* and *BMPER* in skeletal tissues during craniofacial development, to better understand the role of *BMPER* as an evolutionarily conserved BMP regulator, and to clarify the contribution of both genes to the genetic risk for CS. Using an assay for enhancer activity in transgenic zebrafish, we examined candidate sequences in the intergenic region between *BBS9* and *BMPER* and the intronic region of *BBS9* identified as a CS risk locus. We identified two sequences within the risk locus near *BMP2* that regulates expression during craniofacial development. We similarly found multiple enhancers intronic to *BBS9* and in the intergenic region with activity consistent with endogenous *bmper* expression, including two that are active in skeletal tissues, and one of which is specifically active near the site of frontal bone initiation. Importantly, from each identified risk locus, one of the active enhancers encompasses an SNP associated with CS risk (4). Finally, we identified candidate interactions with transcription factors for the skeletal enhancers from both loci through a yeast one-hybrid assay (eY1H). Therefore, *in vivo* interactions suggest regulatory interactions for future examination in determining the mechanism underlying the genetic risk of CS associated with each regulatory domain.

## Results

### Selection of enhancers and screen results

We examined a 1.3 million base pair (bp) region (chr7:33,197,564-34,501,768) encompassing *BMPER* on GRCh38/hg38, to select sequences to test as regulatory elements. This region includes the risk locus in *BBS9* identified from the GWAS for sagittal NCS (4) and the entire intergenic region between *BBS9* and *BMPER* (**Figure 1A**). We also selected sequences in the risk locus downstream of *BMP2*, including one containing rs18843302, which was previously suggested to act as an enhancer (19) (**Figure 1B**). The primary criterion for selection was conservation across species. Within the disease risk loci, we also referenced the SNPs identified in the GWAS (4) as well as cis-regulatory elements predicted from the ENCODE data (20) (21) (22) (**Figure 1A & B**). For the risk locus near *BMPER* (within *BBS9*), only −707*BMPER* contains a significant SNP (rs10254116*), while −687*BMPER*, −684*BMPER*, and −655*BMPER* are each close to an SNP. For the risk locus near *BMP2*, +421BMP2 contains a significant SNP (rs6117669); +382*BMP2*, +402*BMP2*, +460*BMP2*, and +463*BMP2* are each close to a SNP.

**Figure 1.**
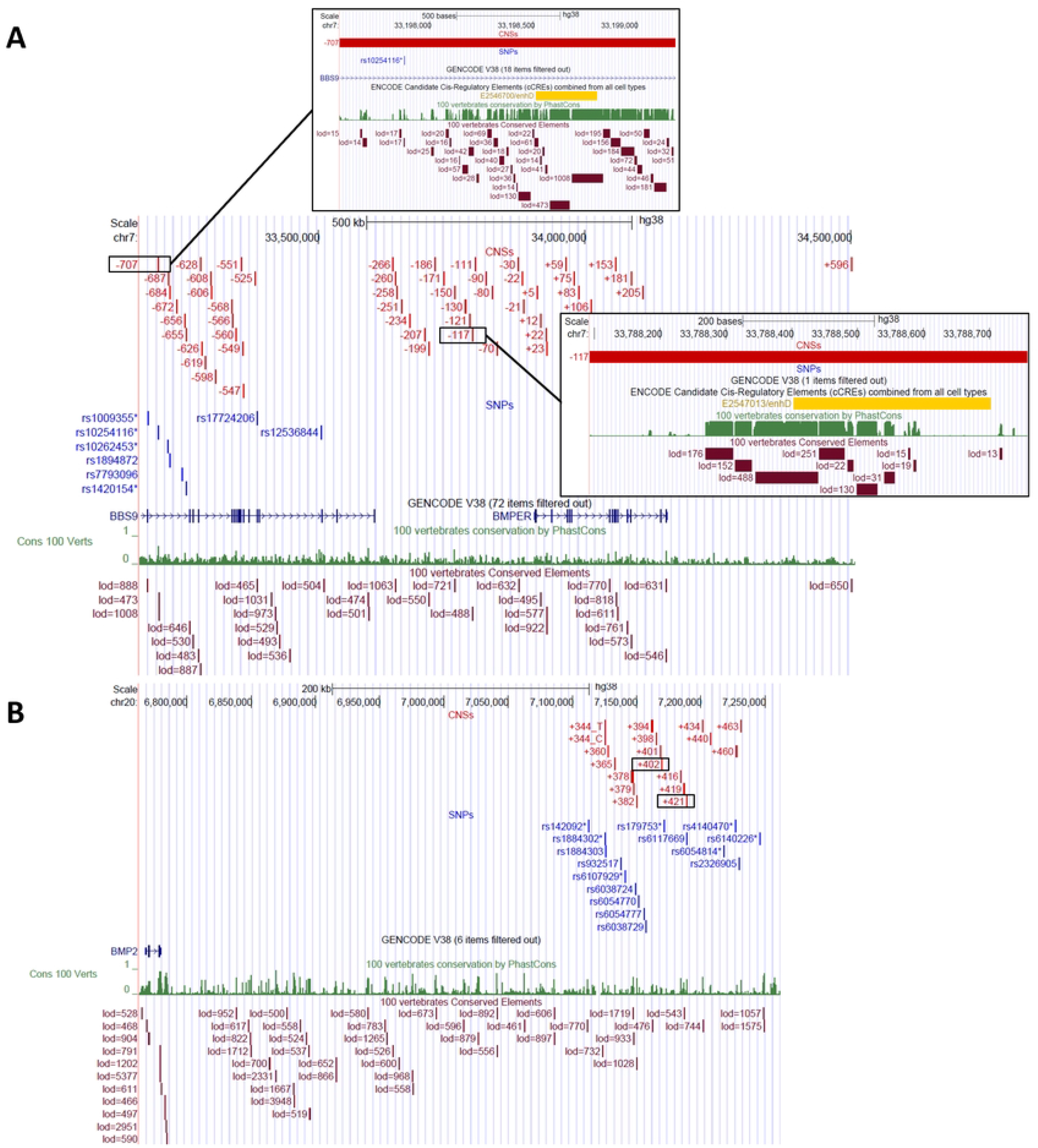
Genomic overview of the genomic regions near *BMPER* and *BMP2*. The custom tracks “CNSs” indicate sequences tested for enhancer activity. SNPs track shows the SNPs significantly associated with sagittal NCS; SNPs* passed the replication study (4). Conservation tracks, PhastCons and Conserved Elements, are the primary criteria used to select sequences. Elements with lod scores above 450 are shown. **(A)** Tested elements for *BMPER*. Boxed sequences are those that showed skeletal-specific enhancer activities in zebrafish transgenesis assay, −707*BMPER* and −117*BMPER*. ENCODE track also indicates predicted enhancer elements for these two sequences. **B).** Tested elements for *BMP2*. Boxed sequences are those that tested positive for skeletal-specific enhancer activities in zebrafish transgenesis assay, +402*BMP2* and +421*BMP2*.

We assessed each candidate element for tissue-specific enhancer activity using zebrafish transgenesis as previously described (23). Each element was cloned into a Tol2-based vector with a minimal promoter and *egfp* (23); embryos injected with each construct were screened for mosaic transgene expression at 5dpf, when the first cranial bone and cartilage elements have formed. To aid in screening for expression in cranial skeletal tissues, we injected into embryos transgenic for *sp7: cherry*, expressed in early osteoblasts. Candidate elements that gave rise to tissue-specific activity were summarized in **Supplemental Tables 1 & 2**. For expression patterns of interest, injected embryos were raised and bred to established transgenic lines. Several sequences for *BMPER* give rise to transgene expression consistent with the endogenous zebrafish *bmper* expression in the skeletal tissues, including pectoral fins, cranial cartilages, and bones (8) (15). Some also showed transgene expression in the inner ear (24) (8) (**Supplemental Table 1**).

For *BMP2*, the enhancer activities consistent with the endogenous *bmp2* genes include pectoral fins, cleithrum (25) (26), otic vesicles (25), and gills (27). The mosaic transgene expressions driven by +372*BMP2* in the notochord and by +382*BMP2* in the gills were particularly specific and strong (**Supplemental Table 2**). The reference SNP near *BMP2* associated with CS risk, rs1884302, had previously been assessed for effect on regulatory function in a similar zebrafish assay (19). We tested the same sequence encompassing the SNP in our assay, comparing the reference and alternate alleles, but we found no tissue-specific enhancer activity above background for either allele. We further characterized the enhancers for each gene with activity primarily in cranial skeletal tissues, −707*BMPER*, −117*BMPER*, +402*BMP2*, and +421*BMP2*.

### −117*BMPER* is a conserved enhancer active in early osteoblasts

From the intergenic region between *BMPER* and *BBS9*, the sequence −117*BMPER* showed enhancer activity in the cranial bones and cartilage of mosaic fish at 5dpf (**Figure 2A-B**). This enhancer has a maximum log odd score of 488 in PhastCons and is conserved down to chicken. It also contains an enhancer element predicted by the ENCODE data (20) (21) (22) (**Figure 1A**). In 5dpf transgenic larvae, *egfp* is expressed widely in cranial cartilage and bone (**Figure 2C-D**), which is consistent across lines from more than three independent founders. Transgene expression remains prominent in Meckel’s and ceratohyal cartilages throughout the larval skull development. During this period of growth, the expression is also visible but less prominent at palatoquadrate, basibranchial, ethmoid plate, opercles, and subopercles. Notably, transgene expression aligns with the endogenous zebrafish *bmper* expression (15), which strongly suggests the enhancer regulates *BMPER* expression. Interestingly, live confocal imaging revealed that −117*BMPER* directed transgene expression at the osteogenic fronts when the frontal bones began to form, and *egfp* expression precedes *sp7: mCherry* marking the osteoblasts (**Figure 2E-I**). During the rapid growth of the frontal and parietal bones, the *egfp* expression is prominent at the osteogenic fronts in the region of osteoblast precursors. The expression is also visible at lateral ethmoids and supraorbital bones (**Figure 2J-L**). This strong expression was transient and became less pronounced as the skull growth slowed and sutures formed at 8-9mm standard length (SL). Therefore, −117*BMPER* most likely regulates *BMPER* expression during early osteoblast differentiation. Notably, after the enhancer activity diminished at the osteogenic fronts, it became prominent at the supraorbital lateral line canals (data not shown), another site of rapid bone growth and remodeling.

**Figure 2.**
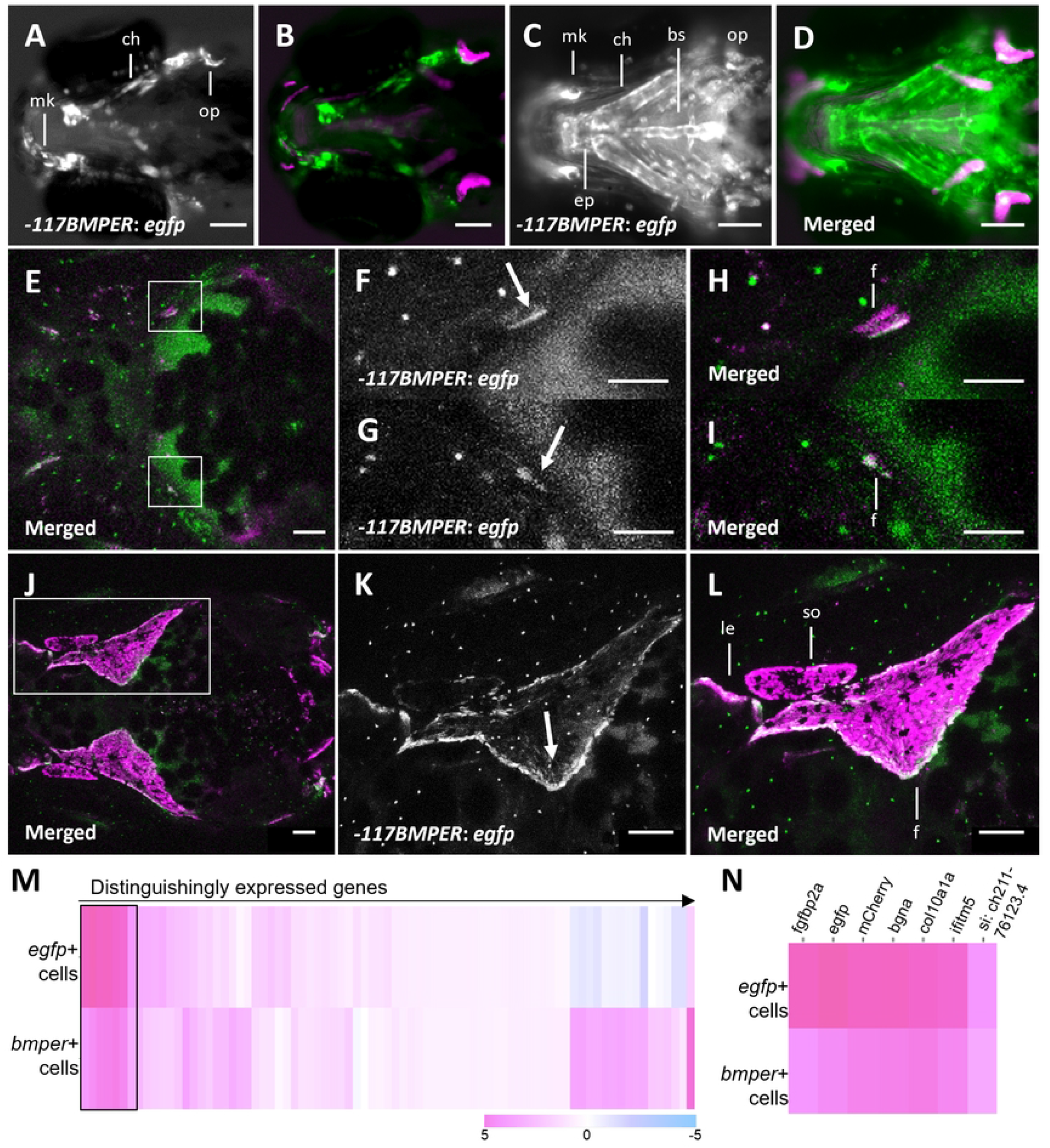
-117*BMPER* directs *egfp* expression to developing craniofacial bones and cartilage in zebrafish. **A-B)** Ventral view of 5dpf zebrafish mosaic for −117*BMPER*. A) Enhancer is active in early chondrocytes and osteoblasts. B) Overlap is shown with osteoblast marker *sp7: mcherry*, in magenta. **C-D)** Ventral view of 5dpf stable transgenic Tg (−117*BMPER: egfp; sp7: mcherry*) zebrafish. Enhancer is broadly active in craniofacial bones and cartilage. **E-I)** Compressed Z-stacks from confocal imaging of a transgenic Tg (−117*BMPER: egfp*; *sp7: mcherry*) animal at 5.2mm SL (19dpf). Enhancer activity is prominent at the osteogenic fronts of both developing frontal bones (arrow). **J-L)** Dorsal view of enhancer activity in the same transgenic animal at 6.62mm SL (26dpf); each image is a maximum intensity projection of confocal slices. Enhancer is active in early differentiated osteoblasts, the arrow points to the osteogenic front and the direction of growth. **M-N)** ScRNAseq was performed on cells isolated from transgenic zebrafish skulls at 2 and 3 weeks. The enhancer, indicated by *egfp* expression, reflects the part of *bmper* expression in osteoblast precursor cells. M) Heatmap of genes (in columns) that are distinguishingly expressed in *egfp+* (first row) and *bmper* expressing osteoblasts (second row). The quantification represents the Log2 fold changes in the expression of each gene in each column relative to the entire dataset. N) Significantly upregulated genes based on adjusted P-value (P <0.05), in either *egfp+* cells or *bmper+* cells, boxed in M). The transgene *mCherry* directed by *sp7* marks osteoblasts. bs: basibranchial; ch: ceratohyal; ep: ethmoid plate; f: frontal bone; le: lateral ethmoid; mk: Meckel’s cartilage; op: opercle; so: supraorbital. Scale bars are 100μm in A-E, J-L, and 50μm in F-I.

The imaging data show the activity of −117*BMPER* in subsets of cartilage and bone cells throughout skull development. To further characterize the specificity of enhancer activity, we performed single-cell RNA sequencing (scRNAseq) on dissected skulls of 2–3-week-old fish transgenic for −*117BMPER: egfp* and *sp7:mcherry*. We generated a heatmap of distinguishingly expressed genes, comparing *egfp* and *bmper* expressing cells (**Figure 2M-N**). The expression level for each gene is indicated by the log2 fold changes of expression in each cell group relative to other cell clusters. Consistent with −117*BMPER* acting as an enhancer for *BMPER, egfp* is significantly upregulated in *bmper*-expressing cells (**Figure 2N**). Importantly, although the expression profiles of *egfp* and *bmper* expressing cells do not completely overlap, they show the greatest overlap in osteoblasts indicated by the comparable expression level of *col10a1a* (15), *bgna* (28), and *sp7: mCherry* (29) (**Supplemental Table 4**).

### −707*BMPER* contains a risk-associated SNP and is active at the site of frontal bone initiation

-707*BMPER* in the intronic region of *BBS9*, is highly conserved, with a highest log odd score of 1008, and aligns with sequences down to *Xenopus* tropicalis. It also overlaps a predicted enhancer element based on ENCODE data (20) (21) (22) and contains the risk-associated SNP rs10254116 (**Figure 1A**). In mosaic fish at 5dpf, *egfp* is expressed in many major cranial cartilages including ceratohyal and Meckel’s cartilage (**Figure 3A**). We established transgenic lines from three independent founders. Two lines have prominent *egfp* expression in cranial cartilage, similar to the mosaic pattern (−707*BMPER: egfp* (c)), while the third has *egfp* expression predominantly in perichondrium (−707*BMPER: egfp* (pc)). *In-707BMPER: egfp* (c) fish, the enhancer activity is very prominent at ceratohyal, Meckel’s cartilage, and palatoquadrate at 5dpf (**Figure 3B-C**). The ceratohyal expression remains strong until the fish are ~6.15mm SL (~14dpf). As *egfp* expression in the ceratohyal decreases, the expression became visible at the hyomandibula starting around 6.76mm SL (16dpf) and remained prominent through 8.25mm SL (21dpf), encompassing the period when the hyomandibula was ossifying (**Figure 3E-F**). In the perichondral line, the enhancer activity was prominent around Meckel’s cartilage, posterior ceratohyal and palatoquadrate at 5dpf (**Figure 3B and D**). These areas of expression remained strong throughout skull development. In juvenile fish (8.29mm SL, 21dpf), the *egfp* expression co-localized with *sp7: mCherry+* osteoblasts surrounding the ceratohyal cartilage (**Figure 3G-H**). These enhancer activities at ossification sites of cartilage and perichondrium were consistent with the endogenous *bmper* expression in fish at a similar stage documented by Kessels *et al* (16), suggesting −707*BMPER* is an enhancer for *BMPER*. To investigate the enhancer activity of −707*BMPER* at the later stage during skull formation, we performed confocal imaging from both lines during the period of early frontal bone growth (~5-7mm SL). Cells of the zebrafish frontal bone were first detected near the juncture of the taenia marginalis and epiphyseal bar cartilages, and the bone grew on top of the cartilage towards the apex of the skull over the next several days (30). Just prior to frontal bone initiation, −707*BMPER: egfp* (c) fish expressed *egfp* in chondrocytes and perichondral cells in this region (**Figure 3I**), and −707*BMPER: egfp* (pc) fish expressed *egfp* in perichondral cells of the same region (**Figure 3J**). Expression in both lines was maintained as the frontal bone expanded along the epiphyseal bar (**Figure 3 K-P**). Given the evidence that Bmper acts to enhance BMP activity during skeletogenesis, its expression in this area points to a role in positively regulating the early growth of the frontal bone.

**Figure 3.**
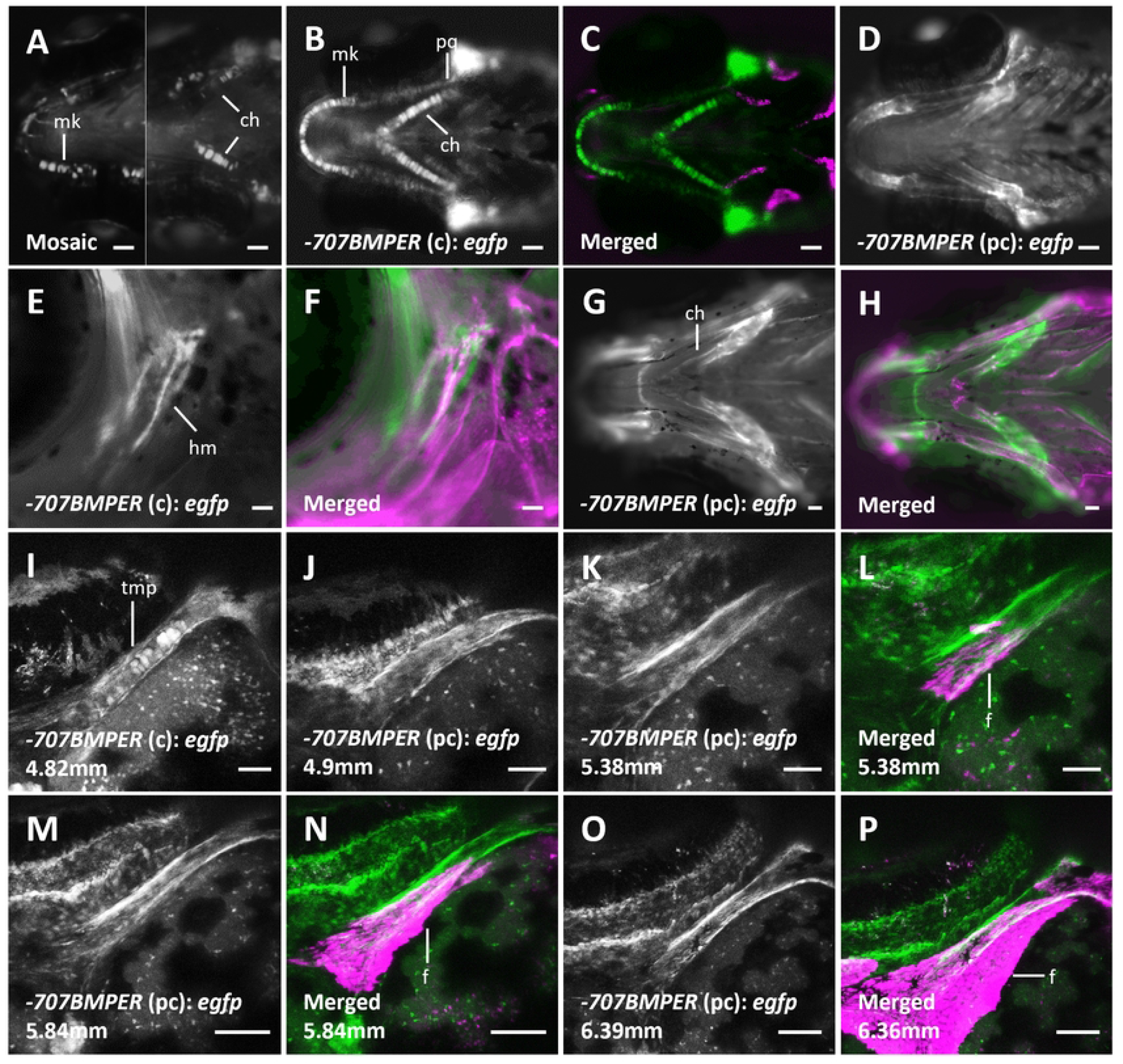
-707*BMPER* directs *egfp* expression to developing craniofacial bones and cartilage in zebrafish. **A)** Ventral view of two 5dpf zebrafish mosaic for −707*BMPER: egfp* expression, showing enhancer activity in early chondrocytes. **B-D)** Ventral view of 5dpf larvae from two independent transgenic lines of Tg (−707*BMPER: egfp; sp7: mcherry*) zebrafish. B-C) In the −707*BMPER*(c): *egfp* line, the enhancer is primarily active in early chondrocytes, with the osteoblast marker shown in magenta; D) In the −707*BMPER*(pc): *egfp* line, the enhancer is prominent in perichondral cells. **E-F)** Side view of a transgenic Tg (−707*BMPER* (c): *egfp*; *sp7: mcherry*) animal at 8.25mm SL (21dpf). At this stage, enhancer activity in cranial cartilage is reduced but is prominent in the hyomandibula. **G-H)** Ventral view of a transgenic Tg (−707*BMPER* (pc): *egfp; sp7: mcherry*) animal at 8.29mm SL (21dpf). Enhancer is still broadly active around craniofacial cartilage, *e.g*. in the posterior ceratohyal. **I-P)** Compressed Z stacks from confocal imaging the two −707*BMPER* transgenic lines during rapid frontal bone growth. I) Dorsal view of the enhancer activity of a transgenic animal of Tg (−707*BMPER* (c): *egfp; sp7: mCherry*) before frontal bone showed up. The enhancer was active at the epiphyseal bar next to the ossification site of the frontal bone. J-P) Dorsal view of the enhancer activity of transgenic animals of Tg (−707*BMPER* (pc): *egfp; sp7: mCherry*). The enhancer was active at the perichondrium of the same part of the epiphyseal bar. ch: ceratohyal; f: frontal bone;; mk: Meckel’s cartilage; pq: palatoquadrate; tmp: taenia marginalis posterior. Scale bars are 50μm in A-F, I-L, 100μm in G-H and M-P.

### +402*BMP2* directs expression in cranial bones

In mosaic fish at 5dpf, +402*BMP2* directed prominent expression in cranial cartilage (data not shown). After germline transmission, transgenic progeny displays broad expression in developing bone. At 5dpf, *egfp* expression completely overlaps with *sp7: mCherry* in branchiostegal ray 1, opercles, maxilla, and cleithrum (**Figure 4A-D**). As the fish grow, *egfp* expression remains primarily on the edges of most cranial bones (**Figure 4E-H**) as well as vertebrae (data not shown). Overall, the enhancer activity of +402*BMP2* is highly specific to bones. There is no predicted enhancer element from ENCODE in this enhancer, but it is 1.7kb away from the CS-associated SNP rs179753.

**Figure 4.**
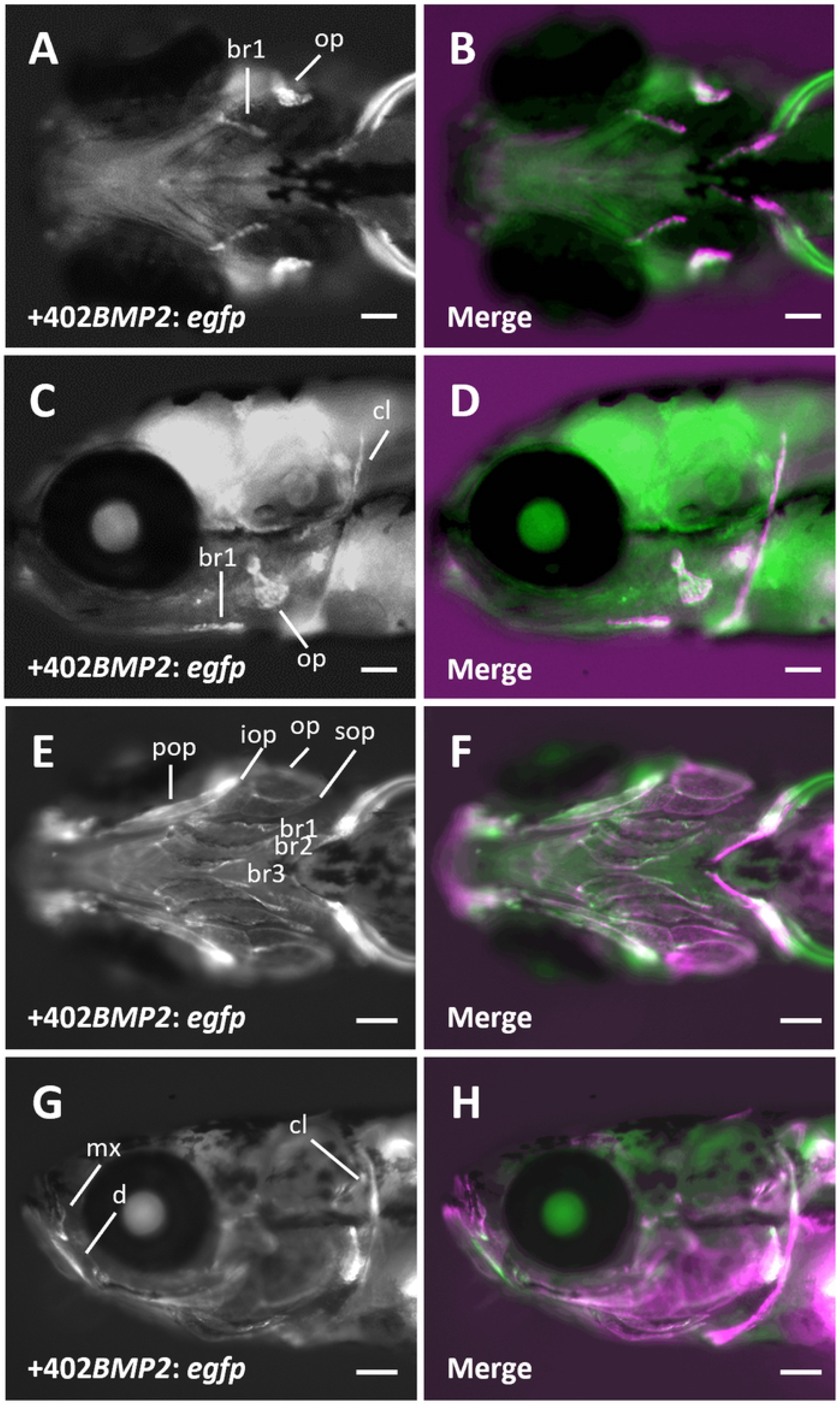
+402*BMP2* directs *egfp* expression to developing craniofacial bones. **A-D)** Ventral and lateral view of a 5dpf stable transgenic Tg (+402*BMP2*: *egfp; sp7: mcherry*) zebrafish. B, D) Overlap is shown with osteoblast marker *sp7: mcherry*, in magenta. Enhancer is active in early craniofacial osteoblasts. **E-H)** Ventral and lateral view of Tg (+402*BMP2*: *egfp; sp7: mcherry*) animal at 6.27mm SL (14dpf). Enhancer is active around the edge of many developing cranial bones, consistent with the position of in early osteoblasts. br 1-3: branchiostegal ray 1-3; cl: cleithrum; d: dentary; iop: interopercle; mx: maxilla; op: opercle; pop: preopercle; sop: subopercle. Scale bar are 100μm in A-D and 300μm in E-H.

### +421*BMP2* directs expression in cranial cartilage

A second enhancer +421*BMP2* within the *BMP2* risk locus has a maximum lod score of 509 and contains the risk-associated SNP rs6117669. There is no predicted enhancer element within +421*BMP2* according to ENCODE. In mosaic larvae at 5dpf, +421*BMP2* primarily directs transgene expression in cranial cartilage. After germline transmission, we isolated a line with broad cartilage expression, including in the ceratohyal, Meckel’s cartilage and pectoral fins (**Figure 5A-B**). At later stages, the enhancer was active in the otic capsule (**Figure 5C-D**) and anterior ceratohyal (**Figure 5E-F**).

**Figure 5.**
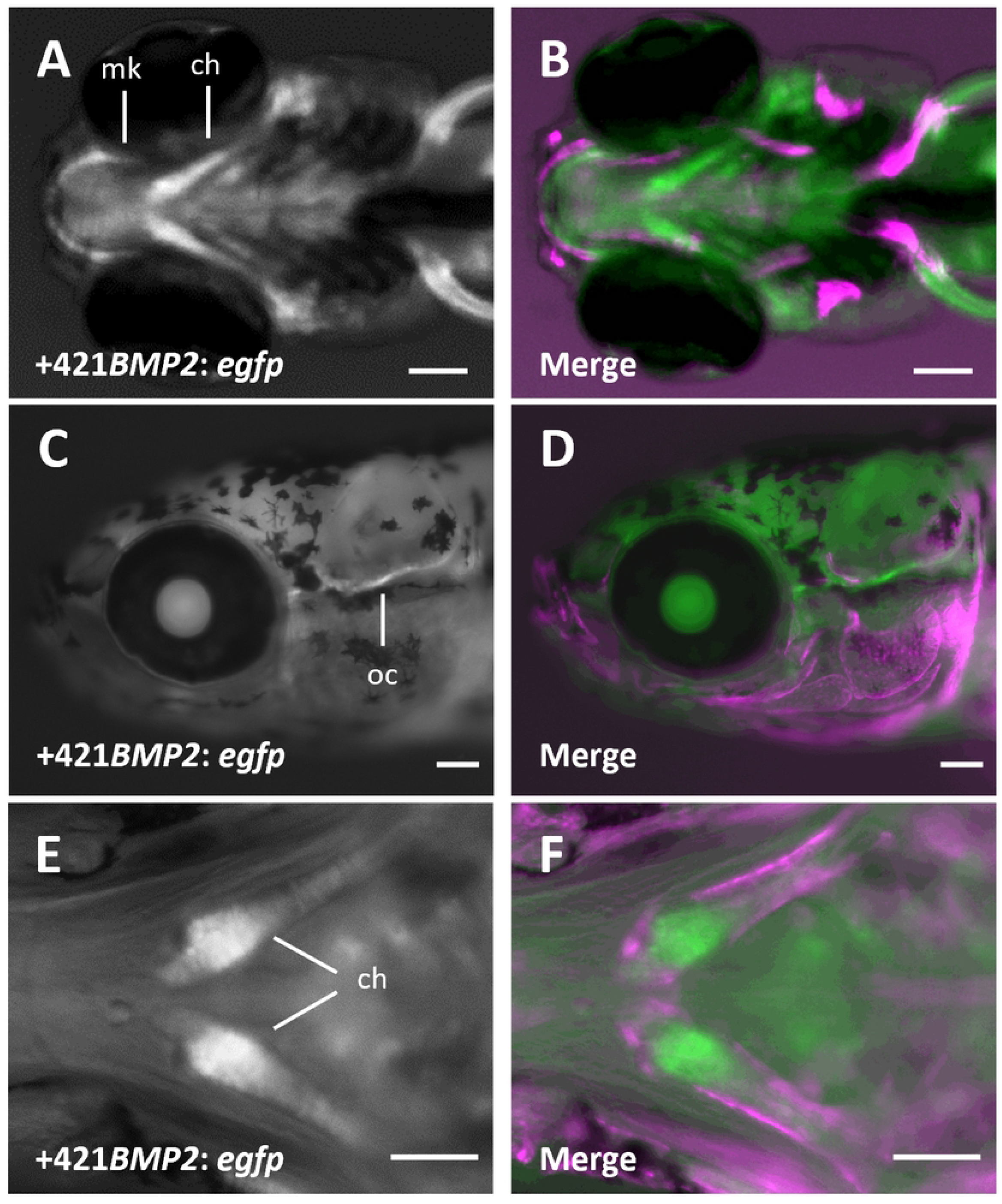
+421*BMP2* directs *egfp* expression to developing craniofacial cartilage. **A-B)** Ventral view of a 5dpf stable transgenic Tg (+421*BMP2*: *egfp; sp7: mcherry*) zebrafish. B) Overlap is shown with osteoblast marker *sp7: mcherry*, in magenta. Enhancer is active in early craniofacial chondrocytes. **C-D)** Lateral view of a stable transgenic Tg (+421*BMP2*: *egfp; sp7: mcherry*) animal at 5.61mm SL (12dpf). Enhancer is active in the otic capsule. **E-F)** Ventral view of a stable transgenic Tg (+421*BMP2*: *egfp; sp7: mcherry*) animal at 6.1mm SL (14dpf). Enhancer is primarily active in the anterior ceratohyal. ch: ceratohyal; mk: Meckel’s cartilage; oc: otic capsule. Scale bars are 100μm.

### Identification of transcription factor binding interactions

To identify potential transcription factor (TF) interactions with each enhancer, we screened them against a library of 1,086 out of ~1,600 annotated human TFs via enhanced yeast one-hybrid (eY1H) assays (31). For comparison, we included two previously identified *RUNX2* enhancers with a specific activity in osteoblasts. Each screen was performed in quadruplicate in two independent yeast strains, and only interactions detected by both yeast strains in at least 2/4 replicates were considered positive interactions (**Figure 6A**). Notably, three out of five enhancers, including −707*BMPER*, interact with ERF, an inhibitory ETS transcription factor that directly binds to ERK1/2 (32). ERK1/2 signaling is activated in CS (33), and haploinsufficiency of ERF causes CS (34).

**Figure 6.**
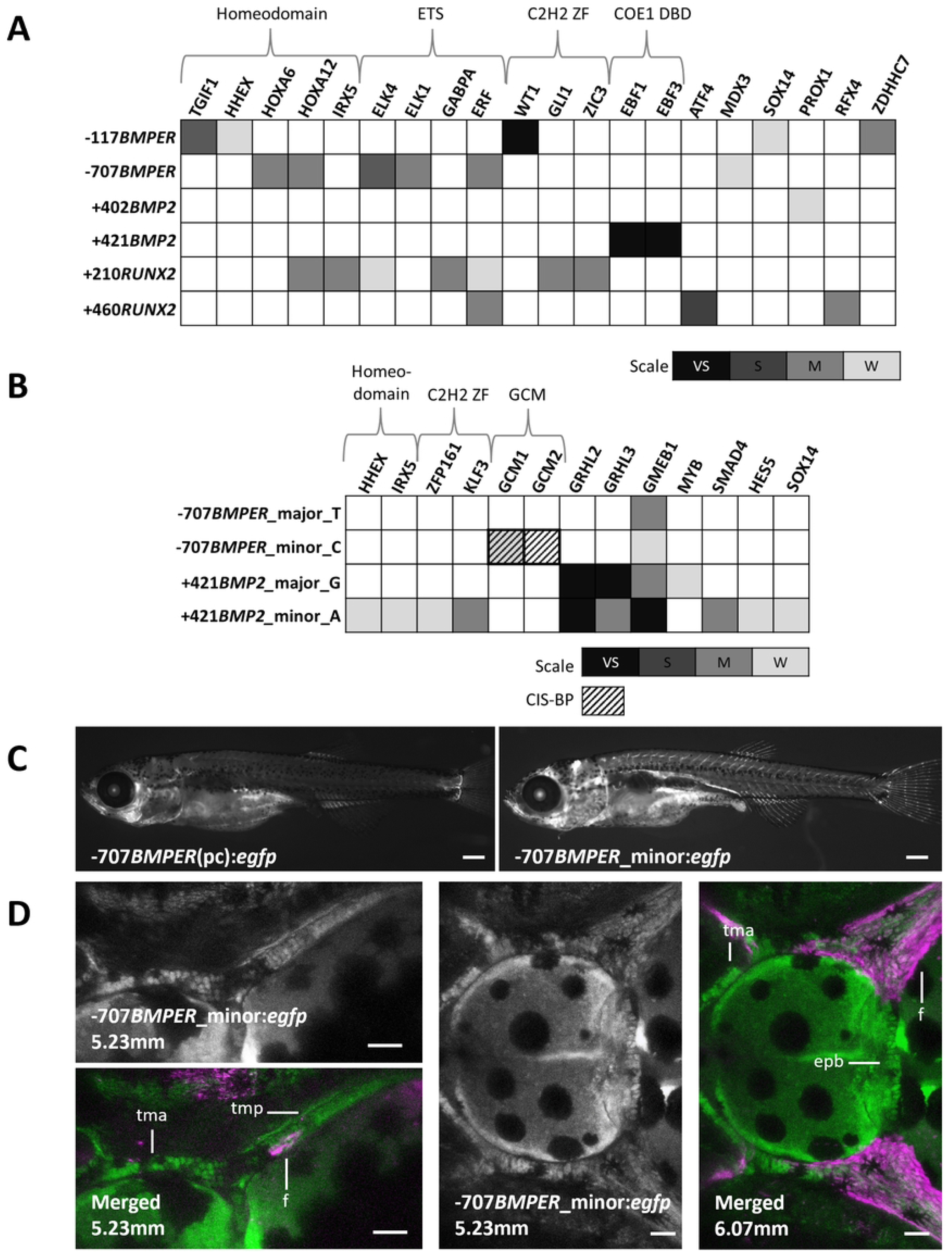
Identification of interacting TFs via eY1H assay and differential enhancer activities of −707*BMPER* by reference and alternate alleles. **A)** TFs that interacted with whole-lengths enhancers. **B)** TFs that interacted with 40bp-sequences around CS-associating SNPs. GCM1 showed up positive in both eY1H assay and CIS-BP differential binding prediction, whereas GCM2 from the same family only did by CIS-BP. The scale indicates the summed strength of TF-bait interactions from two strains of yeasts: very strong (VS), strong (S), medium (M), and weak (W). Some major protein families are grouped based on Pfam database. **C)** Lateral view of −707*BMPER*(pc): egfp with the major allele and −707*BMPER*_minor:*egfp* at 7.1mm (19dpf). The enhancer containing the minor allele showed more extensive activity. **D)** −707*BMPER* enhancer containing the alternate allele drove a broader expression at the site of frontal bone growth. Dorsal view of Tg(−707*BMPER*_minor: *egfp*; *sp7:mCherry*) at 5.23mm and 6.06mm SL. epb: epiphyseal bar; f: frontal bone; tma: taenia marginalis anterior; tmp: taenia marginalis posterior. Scale bar: 50μm.

To test for changes in TF interactions associated with risk alleles, we carried out a targeted screen of the two enhancers containing risk-associated SNPs, −707*BMPER* (rs10254116, C/T) and +421*BMP2* (rs6117669, A/G). For each enhancer, two versions of the 40-base pair sequence centered on the SNP, corresponding to either the reference or alternate allele, were screened as above. In parallel, we used the CIS-BP database (35) to predict differential TF binding around the SNP for each pair of sequences. The results are summarized in **Figure 6B**.

For −707*BMPER*, the alternate allele had a novel interaction with GCM1, one of the two mammalian homologues of the *Drosophila* transcription factor glia cell missing. The interaction was not detected with the reference allele and agreed with the output of CIS-BP, which predicted the binding of GCM1 only with the alternate allele. For +421*BMP2*, there were multiple new interactions both gained and lost for the alternate allele compared to the reference.

We further examined the alternate allele of −707*BMPER* for a functional difference *in vivo*, creating a transgenic line as above for the alternate allele. We found a broader expression, including bones, cartilage, and inner ears, driven by the alternate enhancer in an established transgenic line (**Figure 6C)** (**Supplemental Figure 1**). In particular, in the area of frontal bone initiation, the alternate enhancer drove markedly broader expression that included epiphyseal bar cartilage and cells of the frontal bones in addition to taenia marginalis posterior cartilage seen for the reference allele (**Figure 6D**).

## Discussion

### *BBS9* and *BMPER* constitute a regulatory domain important in craniofacial development

The identification of a risk locus for CS within a *BBS9* intron implicated dysregulation of its expression as a risk factor for CS. However, several lines of evidence suggested an alternative hypothesis that changes in the regulation of *BMPER* underlie the risk. First, CS is not a consistent feature of BBS, and *BBS9* has no known role in osteogenesis. In contrast, *BMPER* has a known role in bone formation, supported by the phenotype of human patients and mouse mutants. Although a null mutation in zebrafish *bmper* has not been described, it is expressed prominently in developing skeletal tissues, suggesting a highly conserved function in skeletal development.

Subsequent to the publication of the GWAS that identified the CS risk locus within *BBS9*, additional support for the existence of broad conserved regulatory domain has been reported. Through comparative genomic analysis of human and chimpanzee cultured cranial neural crest cells, Prescott *et al*. predicted common and differentially active enhancers active in both species. Two of the 5000 most active common enhancers were in introns of *BBS9*, and the entire region encompassing the 3’ end of *BBS9* and the 5’ end of *BMPER* compose a coordinated regulatory region (36). More recently, a pig strain selectively bred for rapid weight gain was shown to carry a 112kb deletion encompassing the 3’end of *Bbs9* and a portion of the intergenic region (37). Obesity is a consistent feature of BBS in humans, and heterozygous loss of *Bbs9* is thought to account for the increased weight gain. However, pigs homozygous for the deletion die in utero. Although the deletion does not affect the *Bmper* gene, *Bmper* expression is greatly decreased in the homozygous fetuses, suggesting loss of critical enhancers that resulted in an intolerable level of Bmper for the proper development and contributed to the lethality. Our identification of two skeletal enhancers, from the intergenic region and from a *BBS9* intron, are also consistent with a large regulatory domain with critical and conserved function in craniofacial development. Importantly, neither of our enhancers was identified by Prescott *et al*.(36), and the addition of −707*BMPER* extends the upstream limit of the regulatory domain to almost 300kb. Finally, *BBS9* and *BMPER* are part of a single topological associated domain (**Supplemental Figure 4**), further supporting a conserved regulatory domain encompassing both genes.

### Presence of skeletal-tissue-specific enhancers within the risk locus near *BMP2*

Justice *et al*. tested a sequence encompassing the lead SNP rs1884302 for enhancer activity using a similar zebrafish transgenesis assay, comparing the reference and alternate alleles. They reported a higher level of expression in mosaic larvae from the alternate allele relative to the reference allele, although the activity was not in skeletal tissues and was only observed in F0 larvae (19). In our assay, we failed to observe any expression above the background driven by either sequence (+344*BMP2* T and C). It is possible that differences in the transgenic vectors can account for our failure to replicate the published results. However, our identification of an additional enhancer, +421*BMP2*, containing a risk–associated SNP and with activity in skeletal tissue suggests that functional changes of this enhancer contribute to disease risk. Of interest, subsequent targeted sequencing of CS patients carrying the *BMP2* linked risk allele showed a linked haplotype of >100kB with 116 significant sequence variants located either in the intergenic region between *BMP2* and the noncoding RNA gene LINC01428 or in the intronic regions of LOC101929265 (38) and the authors suggest that the cumulative effect of more than one sequence variant could underly the increased CS risk. Significantly, an additional enhancer identified in our study with activity in craniofacial cartilage contains one of the newly identified SNPs (+365*BMP2*, **Supplemental Table 3**). Our work provides a functional test for putative enhancer elements in this linked haplotype, which will help define the causal sequence variants in future studies.

### Enhancer-TF interactions suggest underlying regulatory mechanisms

To find regulatory interactions with the identified enhancers and potentially reveal the mechanisms of increased CS risk, we took an unbiased approach to identify TF interactions through an eY1H assay. In contrast to *in silico* predictions, which have a high rate of false positives, TF interactions identified through eY1H show a subsequent verification rate of as high as 40-70% in *in vivo* assays (39) (40) (41). We analyzed the four identified skeletal enhancers and examined previously characterized *RUNX2* enhancers with specific expression in developing bone for comparison. We identified robust interactions implicating multiple signaling pathways, including some already known to play important roles in craniofacial development. These interactions generate specific predictions about developmental regulation that can be tested directly in future experiments to verify their *in vivo* function in regulating skeletal gene expression.

Among the interactions detected, we found that ETS2 repressor factor (ERF) bound to the −*460RUNX2, +210RUNX2*, and −707*BMPER* enhancers. Significantly, haploinsufficiency for *ERF* is associated with cases of both non–syndromic and syndromic CS (34). ERF is an atypical member of the ETS2 TF family that acts as a transcriptional repressor rather than an activator and is thought to prevent the binding of activating ETS factors at the same site. *Erf* and *Runx2* share similar expression patterns at the osteogenic fronts of parietal bones in mice and were shown to act antagonistically in function (34). Our scRNAseq data from skulls of 2- and 3-week-old fish suggests that zebrafish *erf* and *bmper* are likely to be co-expressed in cartilage cells but not in osteoblasts, consistent with the enhancer activity of −707*BMPER* (**Supplemental Figure 2**). Our results support a specific mechanism in which loss of ERF leads directly to the upregulation of *BMPER* and *RUNX2* through decreased binding to their enhancers. More broadly, our results point to the −707*BMPER* enhancer as a critical convergence point for the regulation of craniofacial development, where either sequence variants in the enhancer itself or alterations in the level of the interacting factor ERF leads to changes in *BMPER* expression, misregulation of BMP signaling, and ultimately CS.

Through the eY1H assay, we also found TGFB-induced factor homeobox1 (TGIF1) interacting with −117*BMPER*. TGIF1 belongs to a subgroup of homeobox proteins that are highly conserved transcription regulators. Mutations in *TGIF1* cause holoprosencephaly (HPE), a common human congenital disease that manifests structural brain and craniofacial defects (42). TGIF1 is a co-repressor of the SMAD2-dependent signaling pathway (43), which is proposed as a potential mechanism of *TGIF1*-related HPE. In mouse models, loss of Tgif protein, either Tgif1 or Tgif2 or both, disturbed the Sonic Hedgehog (Shh) activities via dysregulation of SMAD2-Nodal signaling (44). Another disease model includes dysfunction in retinoic acid metabolism, demonstrated in the zebrafish morphants (45), which is also a risk factor for craniosynostosis (46). Although no specific role of TGIF1has been described in osteogenesis, our scRNAseq data showed that *tgif1* and *egfp* driven by −117*BMPER* are expressed in the same subsets of osteoblasts and cartilage (**Supplemental Figure 3**). Future studies can work to uncover the role of TGIF1 in regulating BMP signaling during craniofacial development.

We used two strategies to probe the functional significance of the putative causal SNPs in our identified enhancers. To look for differences in the *in vivo* activity of −707*BMPER*, we made an equivalent transgenic construct and examined both mosaic expression and patterns of activity in established lines. Our results support the more extensive activity of the enhancer containing the alternate allele in skeletal tissues., including at the site of frontal bone initiation. We also carried out targeted eY1H screens on short sequences centered around the putative causal SNPs for both loci, in parallel with *in silico* predictions of TFBSs using CIS-BP (35). For *+421BMP2*, CIS-BP predicted several differences in binding sites for the minor allele, but these did not correspond to the eY1H results. Instead, there were a number of binding interactions gained by the alternate allele and one lost. For −707*BMPER*, the only difference detected in the eY1H assay was the gain of binding by GCM1 to the minor allele, also predicted *in silico* by CIS-BP.

The two mammalian genes *GCM1* and *GCM2* are homologues of the *Drosophila* gene *glial cells missing* (*gcm*), with well-studied roles in trophoblast and parathyroid cell differentiation, respectively. *GCM1* is expressed in a subset of trophoblast cells and is required for normal placenta development (47). Haploinsufficiency for *Gcm1* in mice causes phenotypes resembling preeclampsia (48), while homozygous mutants fail to form functional placenta (49) (50). Because of the early lethality of *Gcm1* mouse mutant, there is limited information on other possible roles. A conditional mouse allele was used to selectively delete *Gcm1* in kidney, demonstrating its important role in recovery from ischemic injury (51). Mammalian *GCM2* is expressed prominently in cells of the parathyroid, and mutations have been demonstrated its requirement for parathyroid development in both mice (52) (53) and humans (54) (55) (56). Interestingly, mice lacking *Gcm2* still have almost normal circulating levels of parathyroid hormone (PTH), apparently due to the upregulation of *Gcm1* and secretion of PTH by cells of the thyroid (52). This suggests significant functional redundancy for the two mammalian genes, which could also mask their roles in other contexts. The zebrafish genome apparently has no *gcm1* orthologue; zebrafish *gcm2* is 82% homologous in amino acids sequences to human *GCM2* and 73% to human *GCM1* (57). Zebrafish *gcm2* is expressed in parathyroid cells, ionocytes (57), and chondrocytes of the pharyngeal arches (58). Antisense morpholino suppression of *gcm2* confirmed a conserved requirement for the zebrafish gene in parathyroid development and further suggested a role in craniofacial cartilage and bone formation (58). More recently, a *gcm2* mutant revealed its important role in maintaining pH in lateral line neuromasts, resulting in hair cell dysfunction (59). However, the craniofacial development of the mutant was not described, and the mutants failed to inflate their swim bladders and did not survive past larval stages. Online expression atlases indicate other areas of expression for both *Gcm* genes in mice, including during craniofacial development, suggesting they may function in other contexts as well. Interestingly, a genome-wide screen for Gcm binding sites during *Drosophila* development uncovered previously undescribed target genes and roles in new tissues (60). They extended the functional assays to mammalian cells and uncovered potentially conserved roles for the mammalian Gcm proteins. Future studies will focus on confirming roles in craniofacial development for specific binding interactions and directly testing the functional importance of the SNPs in contributing to CS risk.

We have identified enhancers from the risk-associated loci near *BMP2* and *BMPER* that regulate expression in the developing craniofacial skeleton and propose that alternate alleles of both lead to an increase in BMP signaling in skeletogenesis. In particular, the increased activity of the alternate allele of −707*BMPER*, including the area of frontal bone initiation and early growth, suggests a specific model for its contribution to CS risk. We hypothesize that in individuals harboring the alternate allele, increased skeletal expression of *BMPER* leads to increased BMP signaling activity and accelerated bone growth during a critical stage of skull formation. In conjunction with other risk factors, genetic or environmental, this increased growth would in turn, increase the likelihood of CS. This hypothesis is consistent with the analysis of zebrafish and mouse models of CS caused by mutations in *TCF12* and *TWIST1* that implicate accelerated growth of the skull bones and the underlying biological mechanisms (61). There is also evidence that −707*BMPER* is negatively regulated by ERF binding, implicating it as part of the mechanism in CS due to *ERF* haploinsufficiency.

## Materials and Methods

### Zebrafish lines

Zebrafish stocks were maintained in the central aquatic facility at Boston University. All animal experiments were approved by the Institutional Animal Care and Use Committee of Boston University (Approval # TR202200000043). Previously reported zebrafish lines used in this study include *sp7: mcherry* and *sp7*−/− (29). We established additional new transgenic lines in the course of these studies: −117*BMPER*: *egfp*, −707*BMPER* (c): *egfp*, −707*BMPER* (pc): *egfp*, +402*BMP2: egfp*, +421*BMP2: egfp*, as described in the text (23).

### Selection of candidate regulatory elements

Within the genomic regions of interest, selection of sequences for analysis as potential regulatory elements was based primarily on multi-species conservation. We selected candidates based on PhastCons (62) (63). included in the 100 Vertebrate Conserved Elements track on the UCSC genome browser. Sequences containing an element with a conservation log odd score of at least 90 were included in our analysis. We also referenced ENCODE Candidate Cis-Regulatory Elements (cCREs) database (20) (22) (21) to check if selected sequences contain any predicted enhancer elements.

### Zebrafish transgenesis assay

Primers were designed to amplify candidate sequences from human genomic DNA using Primer3Plus, with primers on either end located ~200bp outside conserved elements. Adjacent conserved elements were grouped together when convenient, and amplicons ranged from 401 to 1631bp. Primers used for amplification and the genome coordinates of all amplicons are listed in **Supplemental Table 3**. Purified PCR products were cloned into a Gateway^TM^ entry vector using Invitrogen pCR™8/GW/TOPO™ TA Cloning (Invitrogen, catalog # K252002) followed by LR clonase™ reaction (Invitrogen, catalog #11791100) into the *pGW-cFos-EGFP* vectors for analysis (23). All entry vectors were verified by sequencing prior to LR recombination. Tol2 transposase mRNA was transcribed from the plasmid pCS-TP with the mMessage mMachine kit (Qiagen).

We tested the regulatory potential of each candidate sequence through mosaic transgenesis analysis in zebrafish larvae, as previously described (23). For each sequence, at least 150 embryos were injected at the 1-cell stage with 1nL of injection mix containing Tol2 mRNA (final concentration of 35ng/ul), the purified plasmid (final concentration of 15ng/ul), and 1% phenol red for visualization. We examined injected larvae for tissue-specific expression of *egfp* at4-5 days post fertilization (dpf). For sequences with expression patterns of interest, injected larvae were raised and bred to generate transgenic lines.

### Microscopic imaging

For screening and low-magnification imaging, we sedated fish of interest with 4mg/ml Tricaine/MS-222 and mounted them in 3% methylcellulose. We screened injected and transgenic fish for expression by epifluorescence on Olympus MVX10 and captured images using a high-resolution camera. Images were processed using Image J.

We performed serial confocal imaging of live fish as previously described (30). Briefly, each fish of interest is sedated using Tricaine/MS-222 and placed in 3% methylcellulose to measure the standard length. The fish is mounted in 2% low melt agarose, space cleared around the gills for respiration, and the fish covered with fresh Tricaine solution. Stabilized fish are placed onto the stage of the confocal microscope (Leica TCS LSI-III) for image capture, which typically takes less than 10 min., and recovered in fresh tank water. Imaging may be conducted multiple times on the same individual from 15dpf (~5-6mm) to 35dpf (9-11mm). Image stacks were exported and were processed using ImageJ.

### Single cell sequencing

For single-cell profiling of the developing zebrafish skull, we isolated skull tissue from WT and *sp7* mutant fish at ~5.8mm (2 weeks post fertilization) and ~7mm (3wpf). Five fish were included in each sample. To aid in dissection and transcript analysis, fish carried two transgenes, −117*BMPER: egfp* and *sp7: mcherry*. To enrich for skeletal tissues, brains and eyes were removed during dissection. Each sample was then dissociated as described (64). Dissociated cells were immediately loaded onto a 10x Genomics microfluidic chip processed for single cell RNAseq at the BUMC Single-Cell Sequencing Core. Sequencing on Illumina NextSeq was performed to a depth of at least 1,000,000 reads/cell for each library. Cell Ranger software (v.3.1.0, 10x Genomics) was used for barcode recovery, genome alignment (Ensembl GRCz11) and to generate gene-by-cell count matrices with default parameters for each library. Cell clustering and differential gene expression analysis were performed using Loupe Browser 5.0.

### eY1H assay

To explore possible transcription factors that interact with identified enhancers, we performed a gene-centered yeast one-hybrid (eY1H) assay as previously described (31) (65). Briefly, each enhancer, serving as a DNA bait, was cloned into two reporter constructs, expressing either *HIS3* or *LacZ*, and both constructs were integrated into the yeast genome. The yeast with the DNA bait was then mated to a collection of 1,086 yeast strains expressing a TF fused to the *GAL4* activation domain. Binding to a TF activates expression of the reporter genes, allowing the yeast to grow in the absence of histidine and turning blue in the presence of X-gal. Each interaction was tested in quadruplicate, and the strength of the interaction is determined by the intensity of blue product. For the targeted screening that aimed to identify the differential TF binding at the CS-associated SNP, a 40bp-sequence with the SNP in the middle was used as a DNA bait. The results of targeted screening were also compared with the differential TF binding prediction using publicly available CIS-BP database (35).

## Acknowledgement

This work was funded by National Institutes of Health grants R01 HG005039, U01 DE024434 and R21 DE029916 to SF and R35 GM128625 to JIFB. We thank BUMC Single-Cell Sequencing Core for their support of the single cell sequencing experiments and the staff of the BUMC Aquatics Facility for their expert care of our zebrafish colony. We thank Alexandra M Scalici and Kenta Kawasaki for their guidance and assistance with major experimental techniques. We also thank Kelly Ziyi Miao for her support with experiments as well as offering constructive discussions.

---

**Supplemental Figure 1.**
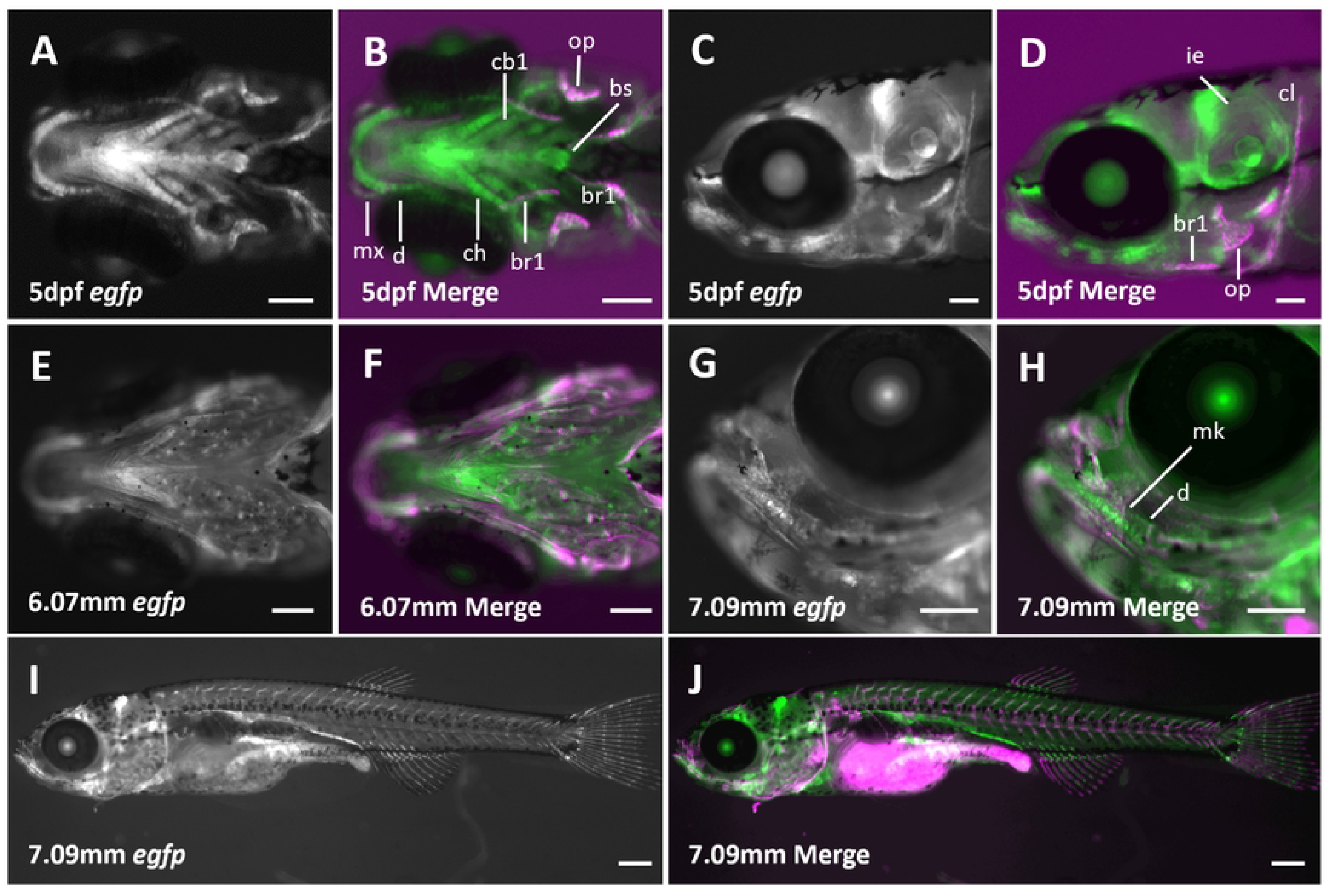

**Supplemental Figure 2.**
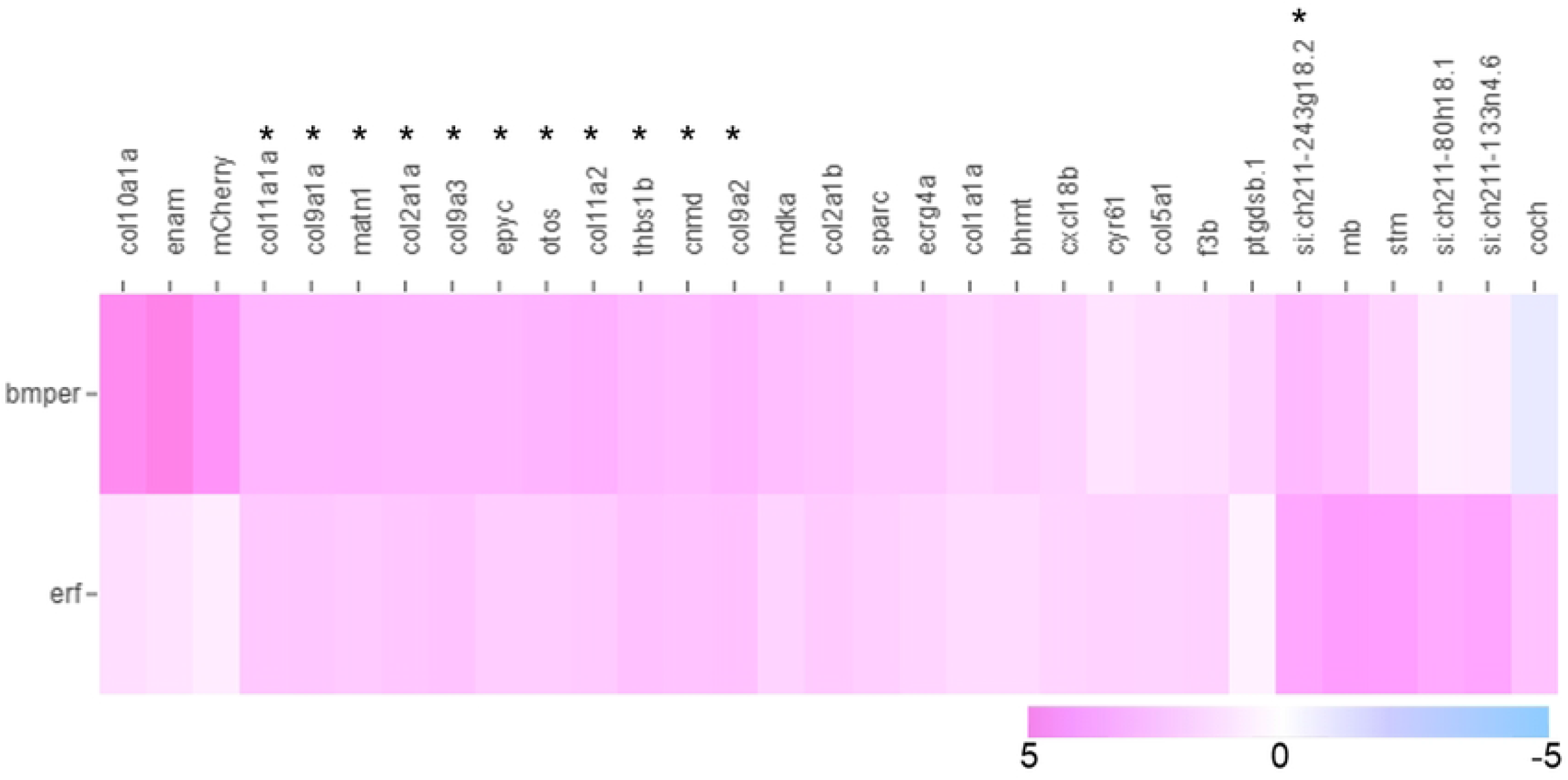

**Supplemental Figure 3.**
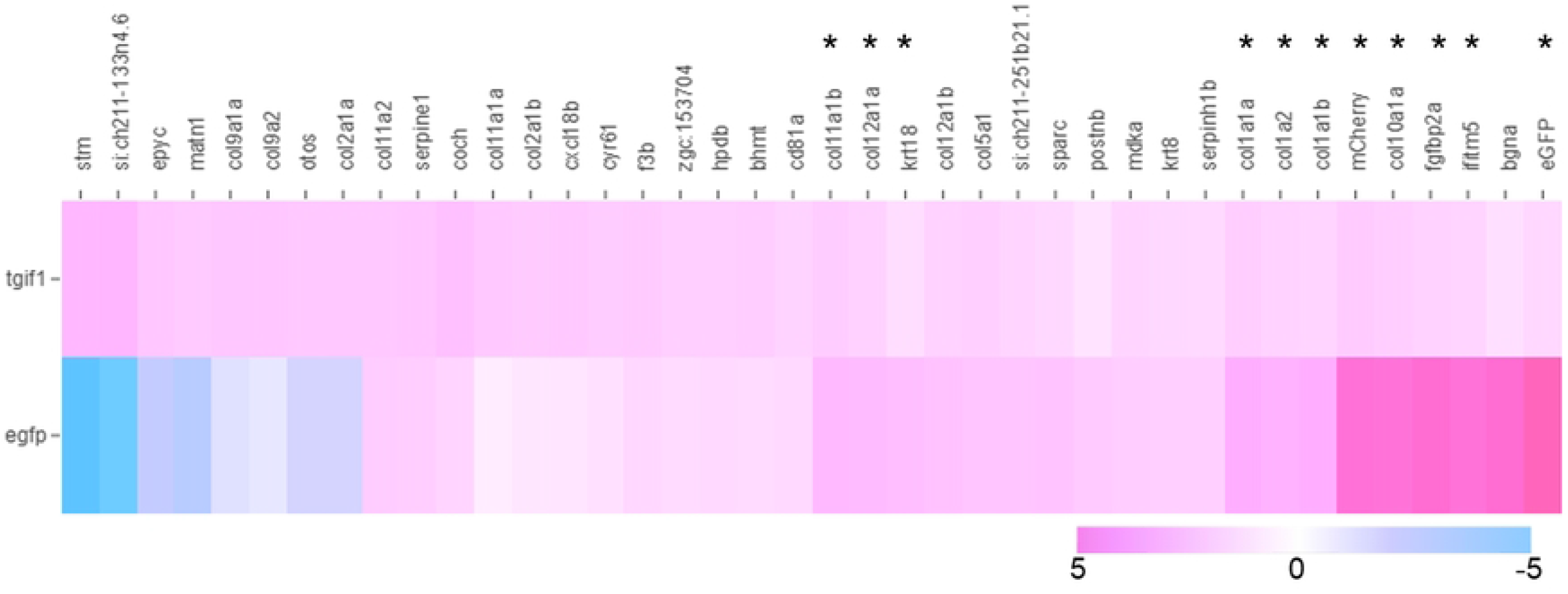

**Supplemental Figure 4.**
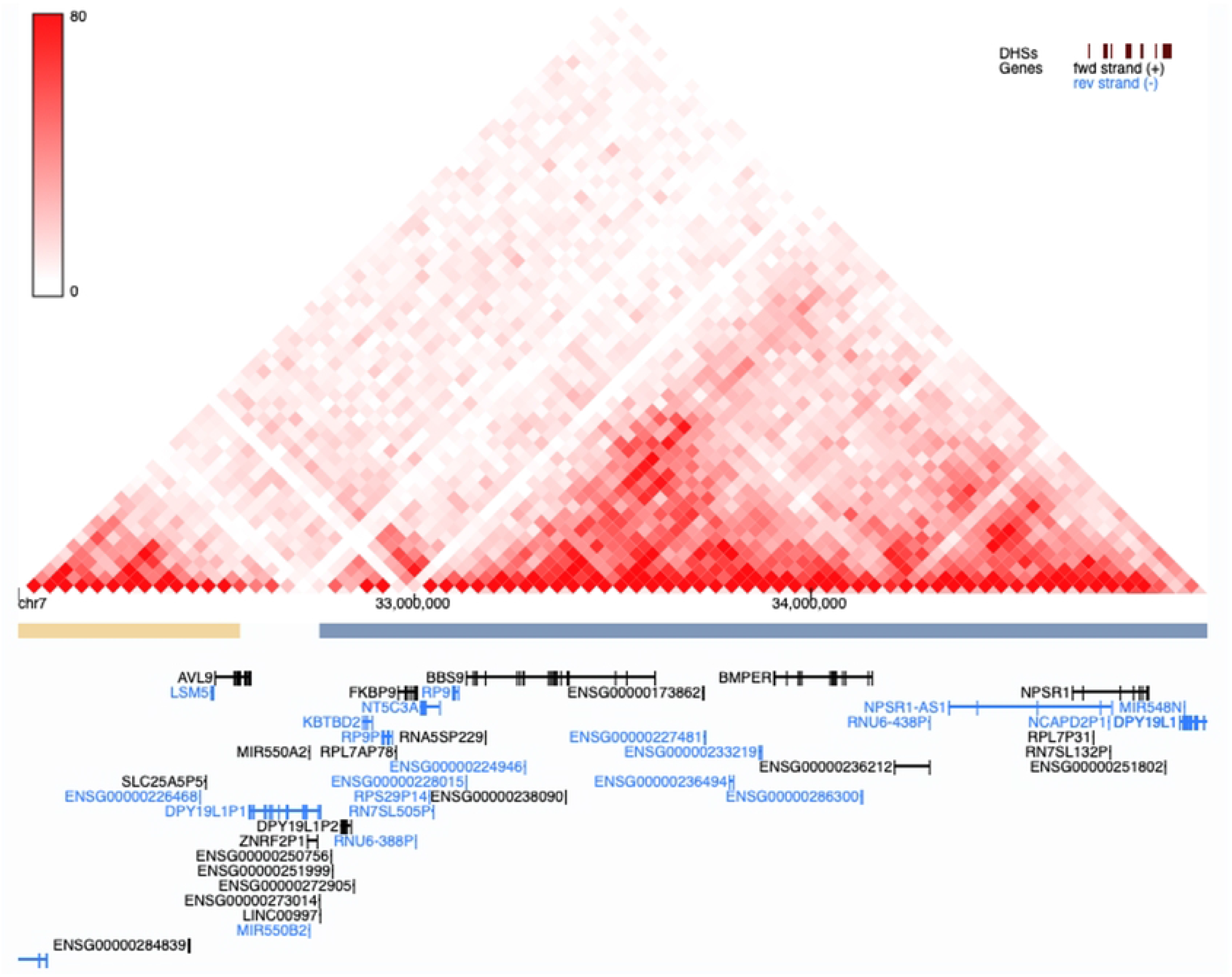

## Notes

### Competing Interest Statement

The authors have declared no competing interest.

